# Revealing Changes in Linear and Nonlinear Functional Connectivity After Psilocybin and Escitalopram Treatment in Patients with Depression

**DOI:** 10.1101/2025.03.05.641592

**Authors:** S.K.L. Quah, C. Glick, L. Roseman, L. Pasquini, R.L. Carhart-Harris, M. Saggar

## Abstract

Major Depressive Disorder (MDD) is typically characterized by altered linear functional connectivity (FC) across large-scale brain networks. Yet, it is unclear whether similar alterations are observed when nonlinear FC is examined. This study investigated how antidepressant treatment (i.e., psilocybin and escitalopram) modulates both linear FC and nonlinear FC in individuals with MDD. Here, we focused specifically on five key canonical brain networks: the Default Mode Network (DMN), Frontoparietal Network (FPN), Salience Network (SAL), Dorsal Attention Network (DAN), and Ventral Attention Network (VAN). Across both treatments, using resting-state fMRI data, we first compared changes in linear and nonlinear FC between responders and non-responders. Responders exhibited increased linear FC within the VAN and greater nonlinear FC within the DMN and VAN than non-responders. We also observed more between-network linear FC for DMN-DAN and nonlinear FC for DMN-VAN in responders than non-responders. Next, we compared treatments and observed that Psilocybin responders showed greater connectivity between FPN-VAN (linear FC), DMN-VAN (nonlinear FC), and SAL-VAN (nonlinear FC) integration than Escitalopram responders, reflecting enhanced coordination and integration between higher-order networks. Conversely, Escitalopram responders exhibited reduced within-network linear FC within the DMN and SAL and between the DMN and VAN, consistent with a dampening of self-referential and salience processing and altered attentional control. These findings highlight potentially distinct mechanisms of action for psilocybin and escitalopram. Incorporating both linear and nonlinear FC analyses provided a novel characterization of these effects, emphasizing the role of these different interactions in antidepressant response. Future studies should investigate the long-term stability of these network changes and their relationship to clinical outcomes.

## Introduction

Major Depressive Disorder (MDD) is one of the most prevalent and debilitating mental health conditions worldwide, affecting approximately 280 million individuals and leading to significant economic and personal burdens. Despite its prevalence and impact, MDD often remains inadequately addressed by current treatments like SSRIs, which take substantial time in inducing clinically meaningful effects, only achieve remission in 37% of patients^1^, and cause substantial side effects. Recent clinical trials with psilocybin, a primarily serotonin 2A receptor (5HT-2AR) agonist, have highlighted its potential as a novel and rapid-acting therapeutic agent for MDD^2–5^. Psilocybin has demonstrated robust and sustained antidepressant effects after one or two dosing sessions, often in combination with psychotherapy or psychological support. Notably, psilocybin’s antidepressant efficacy extends to patients with treatment-resistant depression (TRD)^6,7^and is characterized by a distinct mechanism of action compared to SSRIs^8^.

Despite these promising outcomes, the neurobiological mechanisms underlying psilocybin’s long-term therapeutic effects remain incompletely understood. Neuroimaging studies of psilocybin’s longitudinal effects on brain function have primarily focused on linear correlation-based functional connectivity (FC) changes^9^and remain mixed, with findings suggesting increased FC between specific nodes (e.g., Anterior Cingulate Cortex (ACC) and Posterior Cingulate Cortex (PCC) nodes of the Default Mode Network (DMN) in MDD)^10^, broader between-network integration (DMN, Salience Network (SN), and Dorsal Attention Network (DAN)) in MDD patients^11^, and DMN-hippocampal alterations in healthy individuals^12^. However, these studies often fail to capture nonlinear changes that may be critical for psilocybin’s sustained effects.

The brain operates across multiple levels of organization, from local circuits to global networks, and these interactions are inherently nonlinear. In MDD, studies using graph theory have shown widespread nonlinear disruptions in brain network topology, including reduced global and local efficiency in key networks like the DMN, somatomotor network (SMN), DAN, and visual network (VIN)^13^. Topological analyses of dynamic FC also revealed that MDD patients spend more time in weakly connected brain states linked to self-focused thinking, with altered interactions in prefrontal, sensorimotor, and cerebellar networks correlating with symptom severity. Multimodal approaches that integrate functional and structural data through nonlinear network fusion highlight disruptions in visual processing, motor, language, and cognition areas^14^. These findings suggest that MDD involves complex, system-wide network dysfunctions, emphasizing the importance of advanced analytical approaches to better understand its neural basis.

Given that psychedelics like psilocybin induce profound shifts in perspective and belief systems, their antidepressant effects may stem from large-scale, nonlinear reorganizations that restore flexible brain network dynamics. Research has shown that reductions in brain network modularity following psilocybin treatment correlate with improvements in depression severity^11^, suggesting that increased global integration of functional networks plays a key role in therapeutic outcomes. Additionally, psilocybin has been associated with more frequent and flexible transitions between network states^15^, reflecting a flattened control energy landscape that facilitates dynamic brain reconfiguration. These findings support the idea that psychedelics promote neuroplasticity and cognitive flexibility by enhancing the brain’s ability to transition between states, potentially counteracting the rigid network dynamics characteristic of depression.

Here, we employed both linear and nonlinear FC analyses to investigate how psilocybin and escitalopram modulate brain network organization in individuals with MDD. We leverage Topological Data Analysis^16,17^(TDA), specifically the Mapper tool^18,19^, to explore the nonlinear spatial reorganization of brain activity associated with antidepressant treatments. Unlike conventional neuroimaging methods that rely on linear pairwise correlations or aggregate group data, TDA enables the analysis of complex, high-dimensional, and individual-level brain networks^20,21^. By applying this novel approach to resting-state fMRI data from MDD patients treated with psilocybin or escitalopram, we aim to provide a deeper understanding of the long-term network changes associated with psilocybin compared to conventional antidepressant treatment.

The central hypothesis of this study is that psilocybin induces a persistent reorganization of brain network dynamics, particularly in 5HT-2AR-rich higher-order networks^22^, leading to greater neural flexibility. However, due to the complexity of these network interactions and the novel application of nonlinear methods, this study is best characterized as exploratory. In addition to investigating treatment effects, we examine differences between responders and non-responders, identifying general connectivity changes associated with clinical improvement. Given that MDD has been associated with hyperconnectivity within the DMN^23^, and dysregulation in frontoparietal^24^(FPN), attentional^25^and SAL^26^networks, we focused on changes in within- and between-network connectivity of these networks. Within- and between-network FC of these networks also reflect the integrity of functional modules and the integration between cognitive, attentional, and self-referential systems. While prior studies have demonstrated acute changes in FC following psilocybin administration^12^, the long-term impact on network organization remains unclear. By applying both linear and nonlinear FC analyses, we aim to provide a comprehensive perspective on how psilocybin and escitalopram affect brain network connectivity. Understanding these network-level effects may not only shed light on psilocybin’s antidepressant mechanism but also help optimize its clinical application for individuals with MDD.

## Methods

### Participant recruitment

This study involves a secondary analysis of data previously collected by Carhart-Harris et al.^2^. Briefly, men and women (age: 18-80 years) with a general practitioner-confirmed diagnosis of unipolar major depressive disorder (MDD) were recruited for the double-blind randomized controlled trial (DB-RCT) through formal trial networks, social media, and other referral sources. Eligibility required a Hamilton Depression Rating Scale (HAM-D) score indicating moderate-to-severe depression (≥17 on the 17-item version with a 0-52 range). The diagnosis of depression and the patient’s medical history were verified through their general physician. Participants discontinued all psychiatric medications prior to the trial, ensuring a minimum washout period of two weeks before initiating trial medication. Additionally, any ongoing psychotherapy was halted at least three weeks before starting the trial treatment. Exclusion criteria included an immediate family or personal history of psychosis, serious health conditions deemed unsuitable for trial participation (as assessed by a physician), history of serious suicide attempts, positive pregnancy test, contraindications for selective serotonin-reuptake inhibitors (SSRIs) or magnetic resonance imaging (MRI), previous use of escitalopram (though prior psilocybin use was permitted), and psychiatric conditions that could impair rapport with mental health caregivers within the trial.

Each participant was assigned two mental health professionals to provide support throughout the trial. This team, consisting of a clinical psychologist, psychiatrist, or psychotherapist alongside an equivalent-grade clinician or trainee, fostered a therapeutic alliance before, during, and after each dosing sessions. Baseline assessments were completed 7–10 days prior to the first trial visit. Additional details of the trial can be found at Carhart-Harris et al^2^.

### Trial design

Twenty-nine patients with MDD were randomly assigned to either the psilocybin (n = 14, 4 female) or escitalopram (n = 15, 4 female) treatment arms. Of these, 15 were classified as treatment responders, showing a ≥50% reduction in Beck Depression Inventory (BDI) scores, while 14 were non-responders with a <50% reduction. Notably, the distribution of responders was uneven across treatment groups, with 10 responders and 4 non-responders in the psilocybin group, and 5 responders and 10 non-responders in the escitalopram group. Due to this imbalance, for the analysis comparing psilocybin and escitalopram responders, we selected participants who exhibited reductions above the median BDI reduction within their respective treatment groups. Each participant underwent an eyes-closed resting-state fMRI scan at baseline (pre-treatment) and at the post-treatment endpoint (Figure 1).

**Figure 1.**
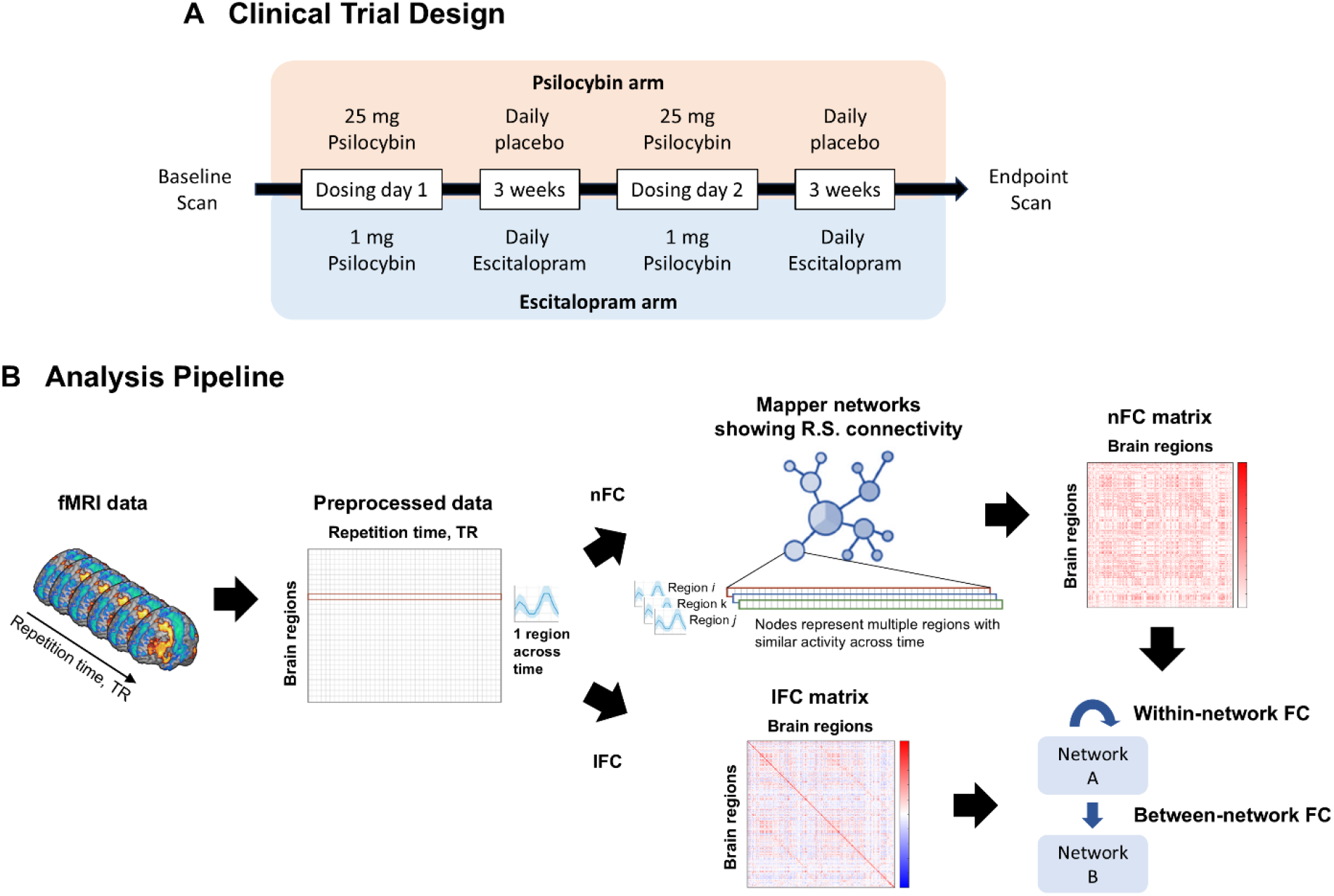
Study design. (a) Participants with MDD underwent a baseline clinical assessment and resting-state fMRI session before being randomly assigned to either the psilocybin or escitalopram group. The psilocybin group received two 25 mg psilocybin dosing days, each followed by three weeks of daily placebo capsules. The escitalopram group received two 1 mg psilocybin sessions, with three weeks of daily escitalopram (10 mg/day after the first session and 20 mg/day after the second). Both groups completed follow-up clinical assessments and fMRI sessions three weeks and one day after the second dosing session. Figure adapted from Daws et. al. 2022. (b) Pre-processed fMRI data is analyzed using two pipelines: the nonlinear FC pipeline, which constructs Mapper networks to capture nonlinear interactions between brain regions, and the linear FC pipeline, which generates linear FC matrices based on Pearson correlation coefficients, reflecting pairwise linear relationships between brain regions. The Mapper networks are then used to compute nonlinear FC matrices by quantifying the topological overlap of network nodes representing different brain regions. Specifically, the strength of nonlinear FC between two regions is determined by the degree to which their respective nodes are co-located within the Mapper-derived low-dimensional manifold. This approach captures nonlinear functional interactions that are not fully characterized by conventional linear FC measures. Finally, within- and between-network FC is derived from these matrices

Dosing day 1 consisted of a 25 mg psilocybin dose (psilocybin arm) or a presumed negligible 1 mg psilocybin dose (escitalopram arm). Participants were informed they would receive psilocybin but were blinded to the actual dosage. A second dosing session (dosing day 2) occurred three weeks after DD1, mirroring the initial dosage. No crossover between doses was conducted. Starting the day after DD1, participants began a six-week course of daily capsules. For the first three weeks, one capsule per day was taken, increasing to two capsules per day for the remaining three weeks. The psilocybin arm received inert placebo capsules (microcrystalline cellulose), while the escitalopram arm received active medication: 10 mg escitalopram daily for the first three weeks, increasing to 20 mg (2 × 10 mg) daily thereafter.

### MRI acquisition

Brain imaging was conducted on a 3T Siemens Tim Trio scanner at Invicro. Eyes-closed resting-state fMRI data were acquired using T2*-weighted echo-planar imaging with 3-mm isotropic voxels. A 32-channel head coil was used to collect 480 volumes over approximately 10 minutes. The imaging parameters included a repetition time (TR) of 1,250 ms, an echo time (TE) of 30 ms, 44 axial slices, a flip angle of 70 degrees, a bandwidth of 2,232 Hz per pixel, and a GRAPPA acceleration factor of 2.256 in-plane field of view, a flip angle of 9 degrees, a bandwidth of 240 Hz per pixel, and GRAPPA acceleration factor of 2.

### fMRI data preprocessing

The fMRIPrep^27^functional preprocessing pipeline begins with generating a skull-stripped reference volume, which is co-registered to the T1-weighted (T1w) image using FLIRT with boundary-based registration. Motion correction is applied using MCFLIRT, and the BOLD time series is resampled to native space. Confound regressors, including framewise displacement (FD), DVARS, and global signals from cerebrospinal fluid (CSF), white matter (WM), and the whole brain, are computed. Motion-contaminated frames are flagged, with additional adjacent frames censored if excessive motion is detected. Denoising involves demeaning, detrending, and regression of nuisance signals, followed by linear interpolation of censored frames and band-pass filtering (0.009–0.08Hz). The final dataset excludes censored frames for further analysis. All activation maps were parcellated into 333 cortical and 14 subcortical brain regions using the Gordon^28^and Harvard-Oxford^29^atlases, respectively.

### Nonlinear FC estimation

The parcellated time-series data (dimensions: brain regions x time frames) were input into the Mapper pipeline following preprocessing. To harmonize data across each brain region, data were z-scored such that each brain region’s time series is standardized. The Mapper pipeline was executed individually for each participant.

The Mapper analysis pipeline involves four main steps. First, high-dimensional input data were embedded into a lower-dimensional space using a nonlinear filter function (classical multidimensional scaling, cMDS) due to its ability to preserve the intrinsic geometric relationships among data points by approximating pairwise distances in the high-dimensional space within a lower-dimensional representation^30^. Second, the lower-dimensional space is divided into overlapping bins for compression and noise reduction. Third, partial clustering is performed within each bin, using the original high-dimensional data to group points into nodes, which helps recover information lost during dimensionality reduction. Finally, a graphical representation of the data is generated by connecting nodes from different bins if they share data points. This process captures the “shape” of the individual’s resting state data to visualize the nonlinear FC of brain regions. Mapper parameters were tuned using an optimization procedure (Saggar et al. X).

Mapper graphs were employed to analyze the topological structure of FC before and after treatment, offering a unique, nonlinear perspective on the organization of brain networks. These graphs represent clusters of brain regions with similar time-series activity as nodes, with edges connecting nodes that share brain regions (Figure 1). This approach captures the “shape” of the FC network and its organization beyond pairwise correlations.

Nonlinear FC between brain regions was determined from the Mapper graphs by calculating the number of shared nodes between each brain region on the mapper graph. FC matrices were calculated for each participant for pre- and post-treatment sessions.

### Linear FC estimation

linear FC was computed using Pearson correlation coefficients between the time-series of brain regions.

### Within-network connectivity

For linear FC, within-network connectivity was measured as the average Pearson correlation between regions within the same functional network. For nonlinear FC, within-network connectivity was defined by the overlap of nodes within a network in the Mapper graph.

### Between-network connectivity

For linear FC, between-network connectivity was defined as the average Pearson correlation between regions in different functional networks. For nonlinear FC, between-network connectivity was based on the overlap between nodes from different networks in the Mapper graph.

### Statistical analysis

To assess differences in FC (FC) changes between responders and non-responders and between responders receiving psilocybin versus responders receiving escitalopram, we conducted a series of independent two-sample t-tests for each network or network pair. To mitigate the influence of extreme values, outliers were identified and removed using the median absolute deviation (MAD) method, which is robust against non-normal distributions. Data points exceeding ±2.5 MAD from the median were excluded from analysis. All statistical tests were two-tailed. Results of all statistical analyses is collated in Supplementary Table 1.

## Results

### Mapper graphs visualizing nonlinear FC

The Mapper-derived insights provide a topological perspective on how nonlinear FC in the brain is altered following psilocybin and escitalopram treatments, complementing traditional correlation-based analyses. Figure 2 illustrates the change in FC matrices and an example of Mapper graphs before and after psilocybin treatment with a random participant, highlighting reorganizations of RSNs.

**Figure 2.**
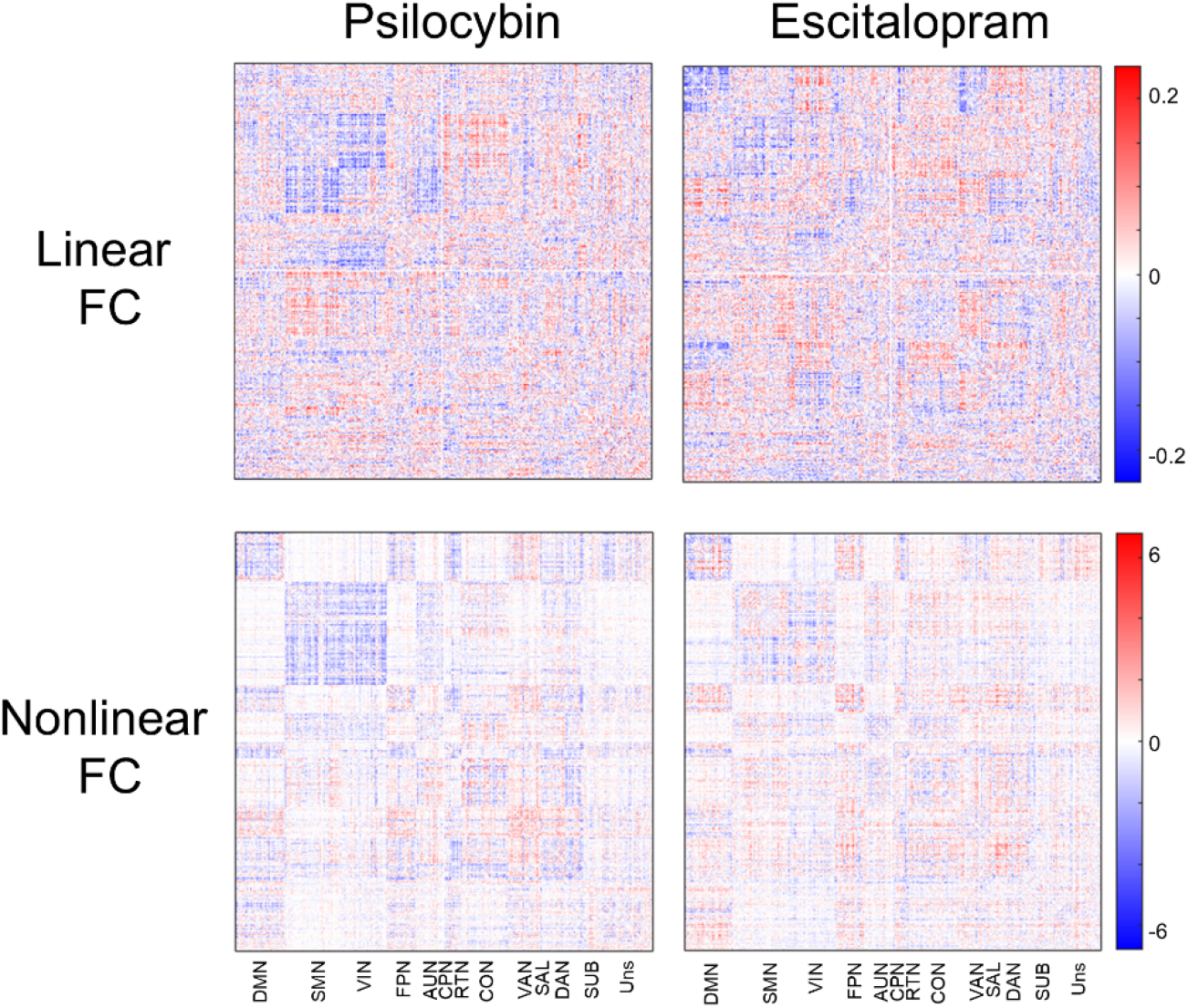
Change in FC matrices after treatment. Participants receiving psilocybin and escitalopram show broad changes in both linear and nonlinear FC. Changes in FC were calculated as the difference between post-treatment and pre-treatment values (post-treatment minus pre-treatment). The DMN, FPN, SAL, DAN, and VAN were the target networks of interest for subsequent analysis investigating within- and between-network FC changes. VIN: Visual Network; VAN: Ventral Attention Network; SUB: Subcortical Network; SAL: Salience Network; SMN: Somatomotor Network; SM-M: Somatomotor-Mouth Network; SM-H: Somatomotor-Hand Network; RTN: Retrosplenial/Temporal Network; FPN: Frontoparietal Network; DAN: Dorsal Attention Network; DMN: Default Mode Network; CPN: Cingulo-Parietal Network; CON: Cingulo-Opercular Network; AUN: Auditory Network. Areas that did not fall into a putative functional network were assigned to “Uns: unspecified”.

### Within- and between-network FC change in responders, combined across treatments

First, we examined changes in within- and between-network linear (linear FC) and nonlinear FC (nonlinear FC) in responders versus non-responders across both treatments (escitalopram and psilocybin combined). (Figure 3).

**Figure 3.**
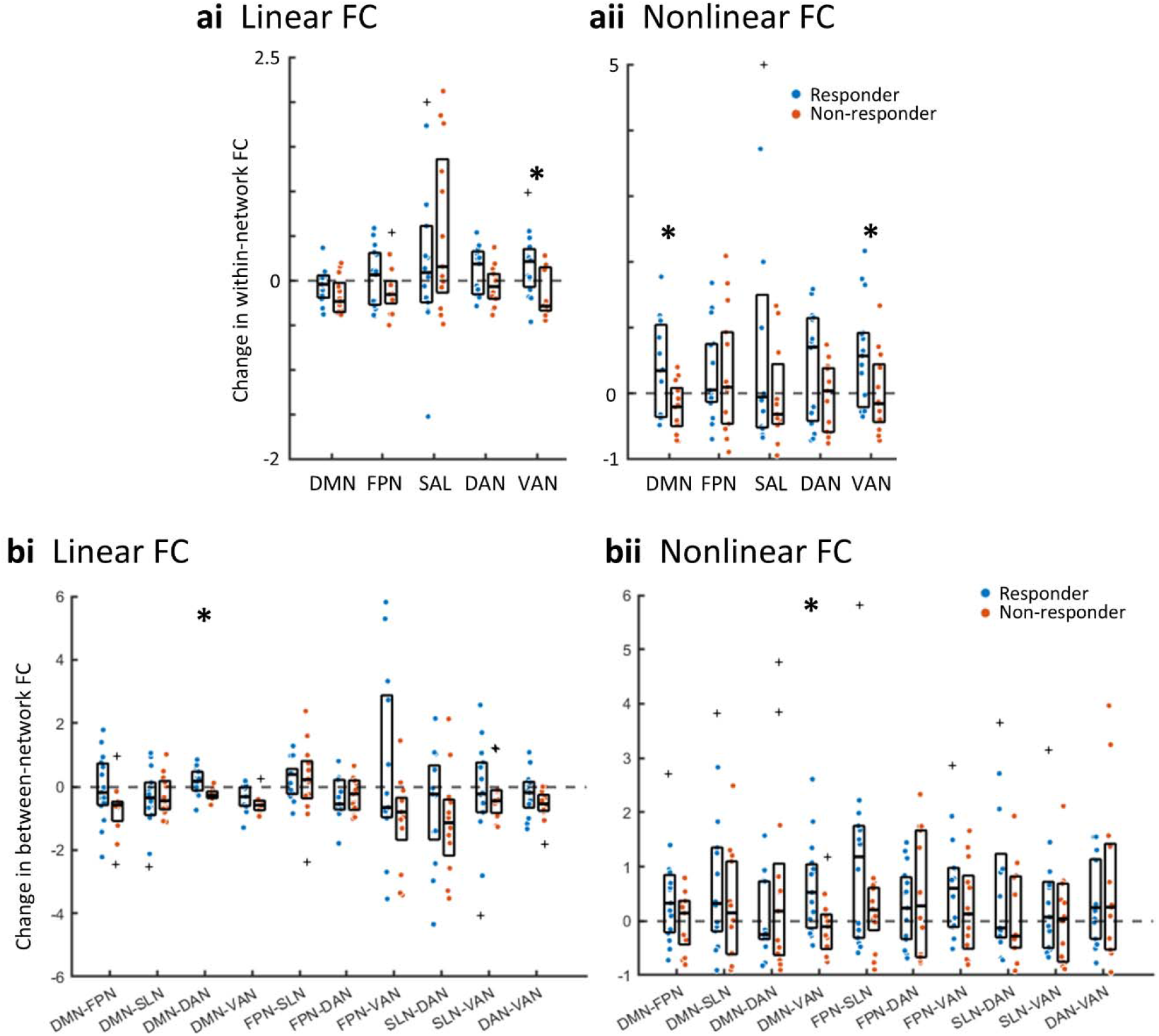
Changes in Linear and Nonlinear Within- and Between-Network FC in Responders. (a) Within-network FC within the VAN was significantly higher in responders compared to non-responders following either treatment. In the DMN, responders exhibited increased within-network connectivity relative to non-responders when assessed using nonlinear FC, whereas no significant differences were observed in linear FC. (b) Comparing changes in between-network FC, responders exhibited increased DMN-DAN FC compared to non-responders, as measured by linear FC. In contrast, nonlinear FC analysis revealed increased DMN-VAN connectivity in responders relative to non-responders. DMN: default mode network; FPN: frontoparietal network; SAL: salience network; DAN: dorsal attention network; VAN: ventral attention network. \**p* < .05. “+” symbols represent outliers.

A significant increase in within-network FC within the VAN was observed in responders compared to non-responders following treatment, as measured by both linear (t = 2.64, p = .014) and nonlinear FC (t = 2.25, p = .033). This converging finding across methods suggests that successful treatment response is associated with strengthened connectivity within this network.

In the DMN, responders exhibited greater within-network connectivity relative to non-responders when assessed using nonlinear FC (t = 2.72, p = .012), but not using linear FC. The selective increase in nonlinear FC within the DMN in responders suggests that nonlinear connectivity patterns within this network may be particularly relevant to treatment response.

These within-network changes were accompanied by alterations in between-network connectivity patterns. Responders exhibited significantly reduced between-network FC between the DMN and DAN compared to non-responders, as measured by linear FC (t = 3.05, p = .006). This reduction suggests that successful treatment response is associated with decreased coupling between the DMN and DAN. NONLINEAR FC analysis further revealed a significant reduction in between-network connectivity between the DMN and VAN in responders compared to non-responders (t = 2.65, p = .014). This suggests that treatment response may involve a decoupling of attentional and self-referential processes in a manner that is detectable primarily through nonlinear connectivity measures, which may capture more dynamic or time-varying functional interactions.

### Comparing within- and between-network FC change between psilocybin and escitalopram responders

Second, we examined changes in within- and between-network FC between responder groups of different treatment conditions using linear and nonlinear FC methods (Figure 4). Among responders, escitalopram treatment was associated with a greater negative change in within-network connectivity within the DMN (t = 2.52, p = .027) and SAL (t = 2.59, p = .027) compared to psilocybin, as measured by linear FC. In contrast, nonlinear FC analyses revealed greater positive change in within-network connectivity within the VAN in responders receiving psilocybin relative to those receiving escitalopram (t = 2.77, p = .016).

**Figure 4.**
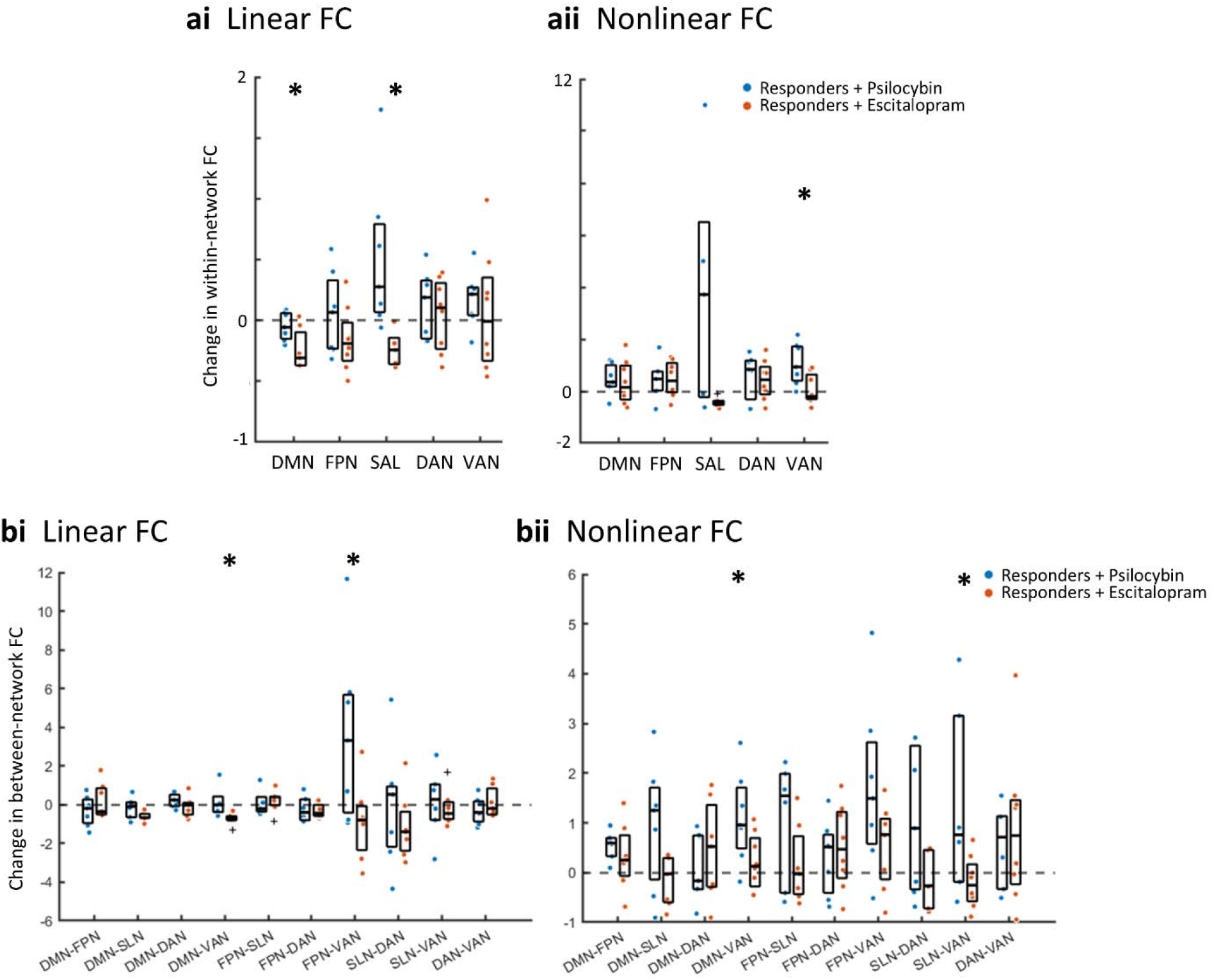
Changes in Linear and Nonlinear Within- and Between-Network FC in Psilocybin and Escitalopram Responders. (a) Among responders, psilocybin treatment was associated with a greater increase in within-network connectivity within the DMN and SAL compared to escitalopram, as measured by linear FC. In contrast, nonlinear FC analyses revealed higher within-network connectivity within the VAN in responders receiving psilocybin relative to those receiving escitalopram. (b) Comparing treatment effects in responders, participants receiving psilocybin exhibited a relative increase in FPN-VAN connectivity as measured by linear FC, and increased DMN-VAN and SAL-VAN connectivity as measured by nonlinear FC compared to escitalopram. Escitalopram responders showed a decrease in DMN-VAN connectivity compared to psilocybin responders. DMN: default mode network; FPN: frontoparietal network; SAL: salience network; DAN: dorsal attention network; VAN: ventral attention network. \**p* < .05.

Linear FC analyses indicated that psilocybin treatment was associated with increased connectivity between the FPN and VAN (t = 2.38, p = .035), suggesting enhanced coordination between cognitive control and attentional networks. Treatment responders receiving psilocybin exhibited a greater increase in DMN-VAN connectivity (t = 2.31, p = .038) and SLN-VAN connectivity (t = 2.20, p = .048) compared to those receiving escitalopram as assessed by nonlinear FC. This finding suggests that effective treatment with psilocybin facilitates greater nonlinear integration between self-referential, salience and attentional networks. Escitalopram responders showed a decrease in DMN-VAN connectivity (t = 2.40, p = .040) relative to psilocybin responders.

## Discussion

This study investigated the effects of psilocybin and escitalopram on brain network reorganization in individuals with MDD, leveraging both linear and nonlinear FC analyses. Our results demonstrate that treatment responders exhibited distinct connectivity patterns compared to non-responders, and that psilocybin responders displayed greater between-network integration within specific higher-order networks relative to escitalopram responders. These findings provide novel insights into the neural mechanisms of antidepressant response and highlight the importance of nonlinear network interactions in understanding treatment effects.

A key finding of this study is that responders exhibited increased within-network FC in the VAN, regardless of treatment type. This suggests enhanced connectivity within this bottom-up attentional network may be a shared neural marker of antidepressant efficacy. The DMN, a core hub implicated in self-referential thought and rumination^31,32^, showed increased within-network nonlinear FC in responders, whereas linear FC analyses failed to detect significant changes. This underscores the importance of nonlinear connectivity methods in capturing network dynamics associated with treatment response.

Additionally, responders demonstrated an increase in between-network connectivity between the DMN and the DAN, as measured by linear FC. Normally, DMN and DAN are anticorrelated^33^, facilitating shifts between internally and externally directed cognitive processes. This coupling is altered in depression^34^. The increase in DMN-DAN connectivity following treatment may reflect a restoration of the normative coupling between these networks. Non-linear FC analyses revealed an increase in DMN-VAN connectivity in responders. As comorbid MDD and anxiety disorder patients show reduced DMN-VAN connectivity^35^, this finding suggests that successful treatment may involve a functional reinstatement of normative connectivity between these networks.

Our exploratory analysis revealed that among responders, psilocybin and escitalopram produced distinct connectivity changes. Escitalopram responders exhibited greater negative change in within-network connectivity in the DMN and SAL relative to psilocybin responders, as measured by linear FC. This suggests that SSRI treatment may work by dampening hyperconnectivity within these networks. Given that increased connectivity within the DMN is a risk factor for developing depression^36^, and the SAL plays a key role in emotional salience detection, a reduction in within-network connectivity within these regions may indicate a normalization of rigid network dynamics, leading to reduced self-focus and emotional reactivity over time.

In contrast, nonlinear FC analyses revealed that psilocybin responders exhibited increased within-network connectivity within the VAN relative to escitalopram responders. The VAN is crucial for bottom-up attentional control and salience detection, dynamically shifting focus toward behaviorally relevant stimuli. Increased VAN connectivity in psilocybin responders may indicate enhanced attentional flexibility, potentially facilitating a more adaptive cognitive state that allows individuals to disengage from rigid, internally focused thought patterns characteristic of MDD.

At the between-network level, psilocybin responders exhibited greater DMN-VAN integration compared to escitalopram responders as detected by nonlinear FC analyses. This finding mirrors the effect seen with responders compared to non-responders. In contrast, linear FC showed that escitalopram responders showed reduced DMN-VAN integration. The DMN-VAN relationship plays a crucial role in dynamically allocating attentional resources between internal and external cognitive states, and increased integration between these networks may underlie the enhanced cognitive flexibility following psilocybin treatment in both MDD patients^10^and healthy individuals^37^. Conversely, the reduced DMN-VAN connectivity in escitalopram responders may reflect greater functional segregation of monitoring internal states and external salience detection. These findings underscore the importance of considering both linear and nonlinear FC analyses when characterizing the differential network-level mechanisms underlying treatment response.

Response to psilocybin treatment also increased connectivity between the FPN and VAN, suggesting enhanced integration of cognitive control and attentional processes, potentially improving cognitive flexibility and adaptive engagement with external stimuli. Additionally, nonlinear FC analyses revealed increased SAN-VAN connectivity, indicating improved salience detection and attentional resource allocation. These findings suggest that psilocybin facilitates a more flexible and adaptive neural state by enhancing interactions across cognitive control, salience, and attentional networks, which may underlie its rapid and sustained antidepressant effects. In contrast, the network changes observed with escitalopram may result from its downregulation of postsynaptic auto-inhibitory serotonin 1A (5-HT1A) receptor in the raphe nuclei^8^. This mechanism contrasts with psilocybin’s ability to broadly modulate neural dynamics, as evidenced by its broad reduction of brain modularity and increased integration between the DMN, SAL and Executive Network (EN) measured using functional cartography in treatment-resistant depressed patients^11^.

While this study provides valuable insights into the neurobiological effects of psilocybin and escitalopram, several limitations should be noted. First, the sample size for imaging analyses was limited, reducing statistical power and preventing correction for multiple comparisons. Given the relatively small sample, we prioritized analyses of higher-order networks and took steps to limit outliers. Future studies with larger cohorts are needed to validate these findings and explore individual differences in treatment response. Second, while we focused on connectivity changes over 6 weeks, the long-term stability of these effects remains unclear. Longitudinal imaging studies could clarify whether network reorganization persists over months and how it relates to clinical outcomes. Additionally, the psychoactive nature of psilocybin presents challenges for blinding, as participants may recognize its subjective effects. Future studies should employ rigorous trial designs to mitigate expectancy biases and explore the relationship between subjective experience and network-level changes. “Another key limitation is the absence of a true placebo control group, as the escitalopram arm received a low-dose (1 mg) psilocybin on dosing days, while the psilocybin arm received placebo instead of escitalopram throughout the trial. This design prevents direct comparisons of psilocybin and escitalopram against a fully inactive baseline, making it difficult to isolate nonspecific treatment effects. Subsequent work incorporating a placebo arm would help isolate drug-specific effects, providing a clearer understanding of how these interventions uniquely modulate brain function in MDD.

In conclusion, this study provides preliminary evidence that psilocybin and escitalopram exert distinct effects on brain network organization, with psilocybin facilitating greater network integration and flexibility, particularly within the DMN and attentional networks. Treatment responders demonstrated increased within-network connectivity in the VAN, a shared neural marker of antidepressant efficacy, while nonlinear FC analyses revealed psilocybin-specific enhancements in DMN-VAN and SLN-VAN integration. These findings suggest that psilocybin’s antidepressant effects may be driven by its ability to disrupt rigid network dynamics^38^and promote neural plasticity^39,40^, contrasting with escitalopram’s more gradual modulation of network connectivity. Importantly, the incorporation of both linear and nonlinear FC analyses allowed for a more comprehensive characterization of these effects, highlighting the unique contributions of different network interactions in mediating treatment response. Future studies should examine the long-term stability of these network changes and their relationship to clinical outcomes to further elucidate the neural mechanisms underlying psilocybin’s rapid and sustained antidepressant effects.

## Reference

1. Warden, D., Rush, A. J., Trivedi, M. H., Fava, M. & Wisniewski, S. R. The STAR*D Project results: a comprehensive review of findings. Curr. Psychiatry Rep. 9, 449–459 (2007).

2. Carhart-Harris, R. et al. Trial of Psilocybin versus Escitalopram for Depression. N. Engl. J. Med. 384, 1402–1411 (2021).

3. Raison, C. L. et al. Single-Dose Psilocybin Treatment for Major Depressive Disorder: A Randomized Clinical Trial. JAMA 330, 843–853 (2023).

4. Davis, A. K. et al. Effects of Psilocybin-Assisted Therapy on Major Depressive Disorder: A Randomized Clinical Trial. JAMA Psychiatry 78, 481–489 (2021).

5. von Rotz, R. et al. Single-dose psilocybin-assisted therapy in major depressive disorder: A placebo-controlled, double-blind, randomised clinical trial. EClinicalMedicine 56, 101809 (2023).

6. Carhart-Harris, R. L. et al. Psilocybin for treatment-resistant depression: fMRI-measured brain mechanisms. Sci. Rep. 7, 13187 (2017).

7. Goodwin, G. M. et al. Single-Dose Psilocybin for a Treatment-Resistant Episode of Major Depression. N. Engl. J. Med. 387, 1637–1648 (2022).

8. Stahl, S. M. Mechanism of action of serotonin selective reuptake inhibitors. Serotonin receptors and pathways mediate therapeutic effects and side effects. J. Affect. Disord. 51, 215–235 (1998).

9. Copa, D. et al. Predicting the outcome of psilocybin treatment for depression from baseline fMRI FC. J. Affect. Disord. 353, 60–69 (2024).

10. Doss, M. K. et al. Psilocybin therapy increases cognitive and neural flexibility in patients with major depressive disorder. Transl. Psychiatry 11, 574 (2021).

11. Daws, R. E. et al. Increased global integration in the brain after psilocybin therapy for depression. Nature Medicine 2022 28:4 28, 844–851 (2022).

12. Siegel, J. S. et al. Psilocybin desynchronizes the human brain. Nature 632, 131–138 (2024).

13. Yang, H. et al. Disrupted intrinsic functional brain topology in patients with major depressive disorder. Mol. Psychiatry 26, 7363–7371 (2021).

14. Chen, N. et al. Estimation of discriminative multimodal brain network connectivity using message-passing-based nonlinear network fusion. IEEE/ACM Trans. Comput. Biol. Bioinform. 20, 2398–2406 (2023).

15. Singleton, S. P. et al. Receptor-informed network control theory links LSD and psilocybin to a flattening of the brain’s control energy landscape. Nature Communications 2022 13:1 13, 1–13 (2022).

16. Munch, E. A User’s Guide to Topological Data Analysis. Journal of Learning Analytics 4, 47–61 (2017).

17. Chazal, F. & Michel, B. An Introduction to Topological Data Analysis: Fundamental and Practical Aspects for Data Scientists. Frontiers Artificial Intelligence Appl. 4, 108 (2021).

18. Geniesse, C., Chowdhury, S. & Saggar, M. NeuMapper: A scalable computational framework for multiscale exploration of the brain’s dynamical organization. Network Neuroscience 6, 467–498 (2022).

19. Haşegan, D., Geniesse, C., Chowdhury, S. & Saggar, M. Deconstructing the Mapper algorithm to extract richer topological and temporal features from functional neuroimaging data. Netw. Neurosci. 8, 1355–1382 (2024).

20. Saggar, M. et al. Towards a new approach to reveal dynamical organization of the brain using topological data analysis. Nature Communications 2018 9:1 9, 1–14 (2018).

21. Saggar, M., Shine, J. M., Liégeois, R., Dosenbach, N. U. F. & Fair, D. Precision dynamical mapping using topological data analysis reveals a hub-like transition state at rest. Nature Communications 2022 13:1 13, 1–19 (2022).

22. Beliveau, V. et al. A High-Resolution In Vivo Atlas of the Human Brain’s Serotonin System. J. Neurosci. (2017) doi:10.1523/JNEUROSCI.2830-16.2017.

23. Hamilton, J. P., Farmer, M., Fogelman, P. & Gotlib, I. H. Depressive rumination, the default-mode network, and the dark matter of clinical neuroscience. Biol. Psychiatry 78, 224–230 (2015).

24. Schultz, D. H. et al. Global connectivity of the fronto-parietal cognitive control network is related to depression symptoms in the general population. Netw. Neurosci. 3, 107–123 (2019).

25. Keller, A. S., Leikauf, J. E., Holt-Gosselin, B., Staveland, B. R. & Williams, L. M. Paying attention to attention in depression. Transl. Psychiatry 9, 279 (2019).

26. Lynch, C. J. et al. Frontostriatal salience network expansion in individuals in depression. Nature 633, 624–633 (2024).

27. Esteban, O. et al. fMRIPrep: a robust preprocessing pipeline for functional MRI. Nat. Methods 16, 111–116 (2019).

28. Gordon, E. M. et al. Precision Functional Mapping of Individual Human Brains. Neuron 95, 791-807.e7 (2017).

29. Desikan, R. S. et al. An automated labeling system for subdividing the human cerebral cortex on MRI scans into gyral based regions of interest. Neuroimage 31, 968–980 (2006).

30. Borg, I. & Groenen, P. J. F. Modern Multidimensional Scaling. (Springer, New York, NY, 2013).

31. Wise, T. et al. Instability of default mode network connectivity in major depression: a two-sample confirmation study. Transl. Psychiatry 7, e1105 (2017).

32. Nejad, A. B., Fossati, P. & Lemogne, C. Self-referential processing, rumination, and cortical midline structures in major depression. Front. Hum. Neurosci. 7, 666 (2013).

33. Fox, M. D. et al. The human brain is intrinsically organized into dynamic, anticorrelated functional networks. Proceedings of the National Academy of Sciences 102, 9673–9678 (2005).

34. Satz, S. et al. The relationship between default mode and dorsal attention networks is associated with depressive disorder diagnosis and the strength of memory representations acquired prior to the resting state scan. Front. Hum. Neurosci. 16, 749767 (2022).

35. Beckmann, F.-E. et al. Specific alterations of resting-state FC in the triple network related to comorbid anxiety in major depressive disorder. Eur. J. Neurosci. 59, 1819–1832 (2024).

36. Posner, J. et al. Increased default mode network connectivity in individuals at high familial risk for depression. Neuropsychopharmacology 41, 1759–1767 (2016).

37. Nayak, S. M. et al. Naturalistic psilocybin use is associated with persisting improvements in mental health and wellbeing: results from a prospective, longitudinal survey. Front. Psychiatry 14, 1199642 (2023).

38. Carhart-Harris, R. L. & Friston, K. J. REBUS and the Anarchic Brain: Toward a Unified Model of the Brain Action of Psychedelics. Pharmacol. Rev. 71, 316–344 (2019).

39. Moliner, R. et al. Psychedelics promote plasticity by directly binding to BDNF receptor TrkB. Nat. Neurosci. 26, 1032–1041 (2023).

40. Zhao, X. et al. Psilocybin promotes neuroplasticity and induces rapid and sustained antidepressant-like effects in mice. J. Psychopharmacol. 38, 489–499 (2024).

